# Unveiling the transcriptome complexity of the High- and Low-Zinc & Iron accumulating Indian wheat (*Triticum aestivum* L.) cultivars

**DOI:** 10.1101/538819

**Authors:** Vinod Kumar Mishra, Saurabh Gupta, Ramesh Chand, Punam Singh Yadav, Satish Kumar Singh, Arun Kumar Joshi, Pritish Kumar Varadwaj

## Abstract

Development of Zinc (Zn), Iron (Fe) and other minerals rich grains along with various stress tolerance and susceptible (STR) wheat genotype, will help to reduce globally spread malnutrition problem. Current study deals with transcriptome profiling of 4 high- and 3 low- Zn & Fe accumulating wheat genotypes (HZFWGs) and (LZFWGs). Functional characterization of expressed and high and low specific genes, accompanied by metabolic pathways analysis reveals, phenylpropanoid biosynthesis, and other associated pathways are mainly participating in plant stress defense mechanism in both genotypes. Chlorophyll synthesis, Zn & Fe binding, metal ion transport, and ATP-Synthase coupled transport mechanism are highly active in HZFWGs while in LZFWGs ribosomal formation, biomolecules binding activities and secondary metabolite biosynthesis. Transcripts accountable for minerals uptake and purine metabolism in HZFWGs are highly enriched. Identified transcripts may be used for marker-assisted selection and breeding to develop minerals rich crops.

## Introduction

Sufficiency of crop production would be the indicator for the livelihood of the farmers and provides essential micronutrients for good health. Micronutrients engage in recreation of the metabolism i.e. the production and functioning of hormones, enzymes, and other substance. Hence, an adequate intake of these micronutrients is necessary for proper growth and development of human and animals [Shenkin, 2006]. The global community is grappling with multiple burdens of malnutrition and 88% of countries are facing a serious burden of either two or three forms of malnutrition. The childhood stunting, anemia in women of reproductive age and/or overweight in adult women are the major problem. More than 40% population of India, mainly women and children are suffering from malnutrition [Global Nutrition Report, 2017]. Zinc (Zn) and Vitamin A deficiency cause a serious problem of diarrhea (children) and multiple micronutrients (Fe and folic acid) supplement required for pregnant woman [Stevens et al. 2013]. Due to severe zinc deficiency approximately 165 million children less than five years of age are stunted, hypogonadism, impaired immune function, skin disorder, cognitive dysfunction and anorexia [Velu et al., 2015]. Fe deficiency leads to decreased immunity, hindered mental development, and higher maternal and prenatal mortality.

Wheat is a major staple food and consumed by 44% world population and major contribution made by developing countries. Minerals concentration in wheat grains and other crops that are grown across Asian are found to be low [Seal et al., 2011; Badigannavar et al., 2016]. Hence, deficiency of Zn & Fe is very common among people especially children and women of South Asian countries. Supplementation of Zn & Fe in diet will be an economic burden for the poor people [Sperotto et al., 2014]. There are few alternative measures *Viz.* fortification by supplementation of minerals in the diet, biofortification, and dietary diversification. Biofortification is an only sustainable method to make available Zn and Fe to the consumer of wheat grain without charging more to the poor people of developing countries [Cakmak, 2008]. Biofortification became possible because of hexaploids spelt wheat (*T. aestivum* var. *spelt*) has a high grain Zn & Fe content and is crossable with bread wheat [Velu et al., 2011; Bouis et al. 2011]. Thus, the spelt wheat is an attractive donor for biofortifying bread wheat. CIMMYT has successfully enhanced the level of Zn & Fe in wheat grain using conventional breeding [Pfeiffer and McClafferty 2007]. There has been continuous progress in terms of identifying competitive high-Zn & Fe candidate varieties in the target countries with the involvement of public and private partners [Velu et al., 2016]. For e.g. two Zn & Fe biofortified varieties (WB02 and HPBW01) have been released in 2017 by Indian Council of Agricultural Research, New Delhi, for commercial cultivation. Initially, to obtain such cultivars multiple time QTLs mapping was performed to developed High- Zn & Fe rich wheat grain development using genotyping and phenotyping of segregating mapping populations [Srinivasa et al., 2014; Velu et al. 2017]. The QTL mapping and identification of polymorphic markers in segregating population is a time-consuming and tedious task with low accuracy [Herrera et al. 2017]. The transcriptome sequencing technology captures the full range of the dynamic spectrum of the transcriptome, with an advantage, when compared to array platforms. Current study deals with the identification of low- and high- Zn & Fe concentration containing wheat seeds. Qualified low and high Zn & Fe containing wheat seeds were sowed in 21 pots having homogenous soil with environmental growth conditions. Further, transcriptome profiling analysis was carried out in 3 replicates leaves samples of 4 high Zn & Fe accumulating wheat genotypes (HZFWGs) and 3 low Zn & Fe accumulating wheat genotypes (LZFWGs), to elucidate three major issues (a) Differences in the transcriptional responses associated between low and high wheat genotypes as well as among them, (b) Identification of different genetic and metabolic changes occurs in accumulation of Zn & Fe in both genotypes (c) To find out associated genes in mineral transportation and responsible for stress tolerance in both types of cultivars. Certainly, this work will be helpful to explore new ways to developed new genotypes of wheat rich in Zn & Fe minerals.

## Results

### Identification of Zn & Fe concentration in used wheat seeds

Zn concentration in leaves of BHU-35, BHU-22, BHU-3, and BHU-24 genotypes were found 47.48 ppm, 47.03 ppm, 45.73 ppm and 44.03ppm respectively, while genotype AM-175, AM-177, and HUW-234 exhibits 38.44ppm, 38.227ppm and 34.358ppm respectively. Moreover, the Fe concentration in all genotypes exhibits in correlation with Zn concentration and found high in HZFWGs as compared to LZFWGs. Interestingly, HZFWGs yield is higher than LZFWGs **(Table 1)** and found that BHU-35 exhibited maximum yield 31.37g/pot followed by BHU-24 (31.19g/pot) and BHU-22 (31.18g/pot) among all genotypes. Furthermore, the significant genetic variation among genotypes was observed, the however same amount of minerals and nutrients was mixed in the soil of each pot.

**Table 1:** Mean performance minerals concentrations and yields analysis of HZFWGs and LZFWGs.

| **Genotype** | **Fe (ppm)** | **Zn (ppm)** | **Yield (g/pot)** |
| --- | --- | --- | --- |
| **BHU-3** | 45.587 | 45.725<sup>H</sup> | 30.385 |
| **BHU-35** | 43.777 | 47.48<sup>H</sup> | 31.3683 |
| **HUW-234** | 38.357 | 34.358<sup>L</sup> | 30.5067 |
| **BHU-22** | 46.745 | 47.033<sup>H</sup> | 31.175 |
| **BHU-24** | 43.267 | 44.03<sup>H</sup> | 31.1933 |
| **AM-175** | 35.535 | 38.44<sup>L</sup> | 26.5517 |
| **AM-177** | 38.265 | 38.227<sup>L</sup> | 30.6 |
| **Mean** | 41.64 | 42.18 | 30.25 |
| **CV** | 5.88 | 5.11 | 3.44 |
| **LSD** | 2.91 | 2.56 | 1.24 |
| **H: High Zinc & Iron content, L: Low Zinc & Iron content** |  |  |  |

### RNA sequencing and transcriptomic profiling of wheat genotypes

Approximately 20.51 to 26.59 million of raw reads with pair-end reads of 100bp were generated for the HZFWGs and LZFWGs samples through RNA sequencing **(Table 2)**. Out of them ∼20.32-26.31 high quality (HQ) reads were observed in all the samples indicating high quality reads generated by Illumina HiSeq 2500. Nucleotide bases in generated HQ read vary from ∼20.52-26.58 billion in seven samples. Then, the HQ reads retained for each sample was used for mapping with recently annotated 1, 37,056 high-quality genes of wheat. The annotated transcripts of each sample with their chromosomal locations, annotated transcript length, COV, FPKM and TPM values and shown in **Supplementary Table S1-S7**. Box plot indicating the level of gene expression (log_2_FPKM) distributed in each sample **(Supplementary Figure S1)**. The genome-wide mapping statistics along with the percentage of mapping for each genotype is given in **Table 2**. The genome-wide mapping of expressed 33767, 29510, 32059, 20152, 35785, 36721, and 29369 transcripts of BHU-22, BHU-3, BHU-24, BHU-35, HUW-234, AM-175, and AM-177 respectively. Expression pattern of transcripts expressed in each wheat genotype is depicted in a circos plot **(Figure 1)**. Venn diagram of HZFWGs and LZFWGs indicates the number of common transcripts among them **(Supplementary Figure S2)**. Moreover, the chromosomal distribution of expressed genes infers that in Chr5D higher numbers of the genes are expressed in each sample **(Figure 2).** Further, the common genes among HZFWGs and LZFWGs were merged and it infers that 2387 and 3933 genes uniquely expressed in HZFWGs and LZFWGs respectively **(Supplementary Table S8-S9)**. The top 450 genes with high FPKM values were selected from HZFWGs and LZFWGs and denoted as High Zn & Fe specific genes (HSGs) and Low Zn & Fe specific genes (LSGs) respectively, listed in **Supplementary Table S10-S11**. Differential gene expression analysis of common 19239 genes from both types of genotype infers that 280 up- and 244 down-regulated genes are expressed **(Figure 1)** and corresponding values are given in **Supplementary Table S12**.

**Figure 1:**
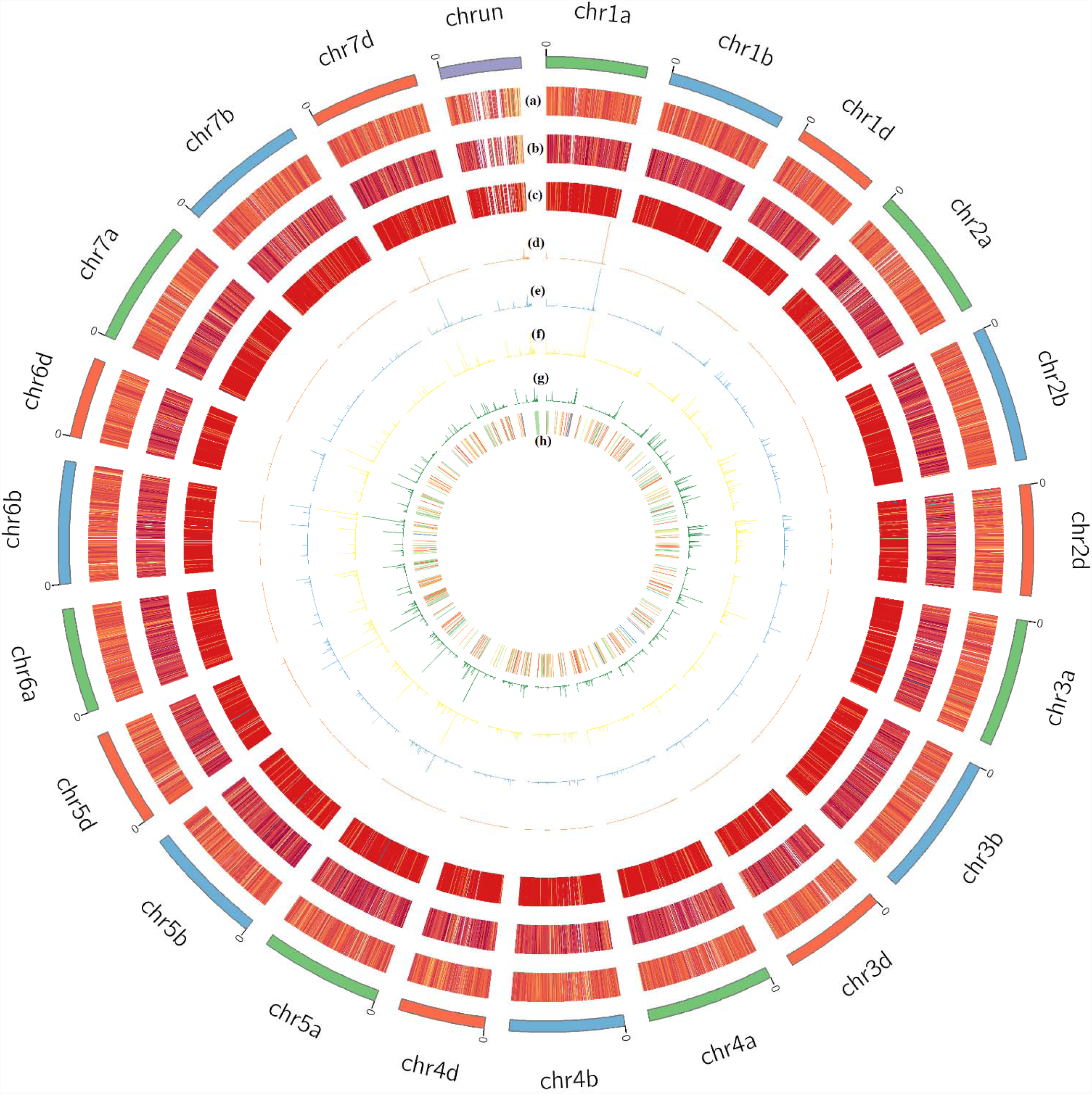
Chromosomal distribution of expressed transcripts for 3 low- and 4 high-Zn & Fe accumulating wheat genotypes in a Circos plot. Heatmaps show the expression values (in different spectral colors) of transcripts wit their gene loci for each low Zn & Fe containing wheat genotypes (a) AM-175, (b)AM-177 and (c) HUW-234. Similarly, line plots show the expression values (high line high expression) of transcripts with their gene loci for each high Zn & Fe containing wheat genotypes (d) BHU-3, (e)BHU-22, (f) BHU-24 (g)BHU-35. (h) Differentially expressed transcripts having fold change values (shown in different spectral color) with their gene locus.

**Figure 2:**
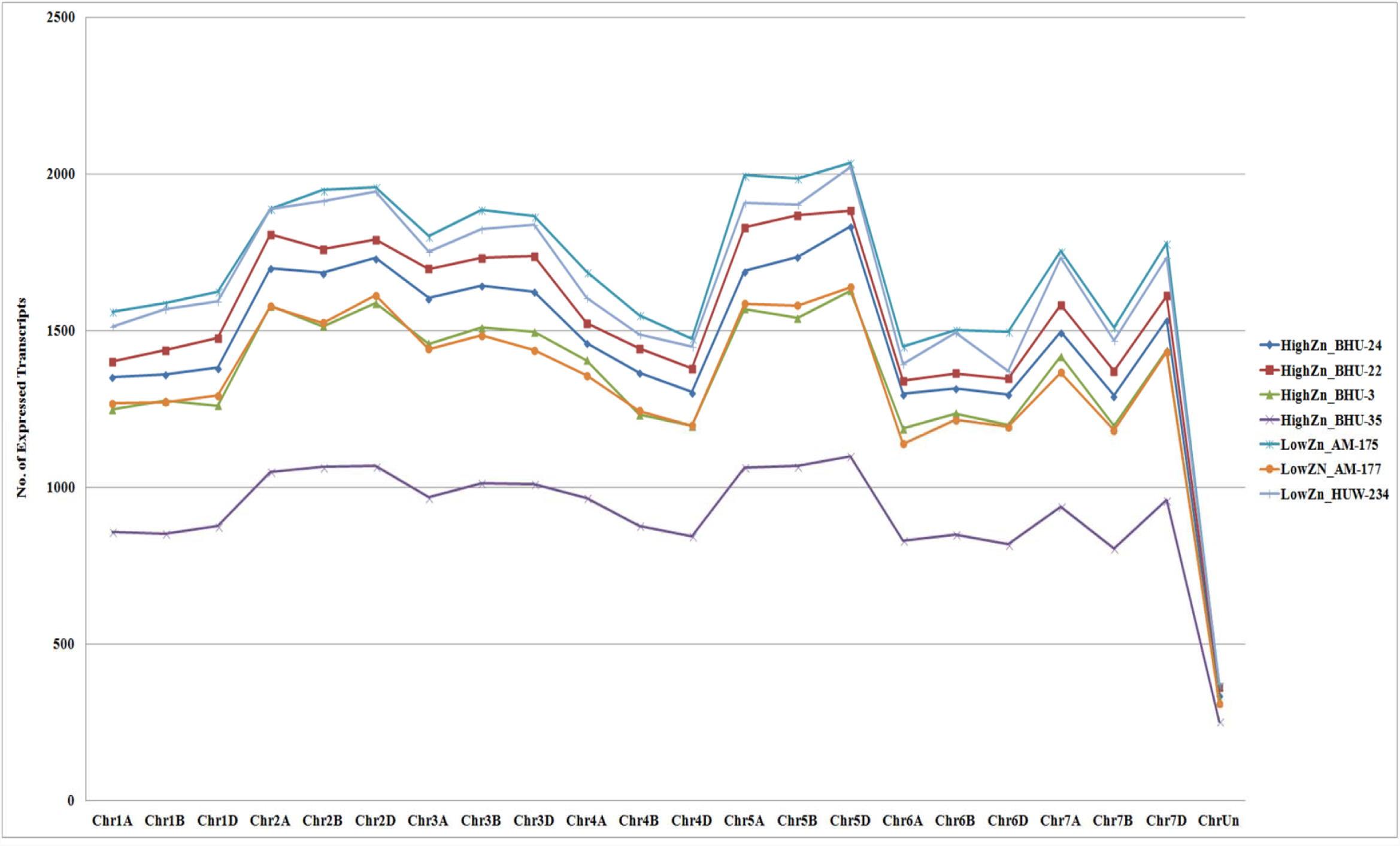
Chromosome-wise distributions of expressed transcripts for each low-and high- Zn & Fe wheat genotypes, denoted as high Zn_ & low Zn_ name of wheat genotype at the bottom of the plot.

**Table 2:** Transcriptome sequencing and de novo assembly statistics of 4 High- and 3 Low- Zn & Fe containing genotypes of wheat.

| Different Statistic /Samples | High Zinc & Iron Wheat Genotypes |  |  |  |  |  |  |  |
| --- | --- | --- | --- | --- | --- | --- | --- | --- |
|  | BHU-22 |  | BHU-3 |  | BHU-24 |  | BHU-35 |  |
|  | BHU22_F | BHU22_R | BHU3_F | BHU3_R | BHU24_F | BHU24_R | BHU35_F | BHU35_R |
| No. of Raw reads | 20513280 | 20513280 | 22182420 | 22182420 | 26598749 | 26598749 | 22952520 | 22952520 |
| No. of HQ reads & Percentage | 20325836<br>(99.09%) | 20325836<br>(99.09%) | 21956299<br>(98.98%) | 21956299<br>(98.98%) | 26319906<br>(98.95) | 26319906<br>(98.95) | 22700054<br>(98.90%) | 22700054<br>(98.90%) |
| No. of bases in HQ Reads | 2052909436 | 2052909436 | 2217586199 | 2217586199 | 2658310506 | 2658310506 | 2292705454 | 2292705454 |
| No. of HQ bases in HQ Reads | 2042078442<br>(99.47%) | 2033874716<br>(99.07%) | 2205663091<br>(99.46%) | 2195544941(9<br>9.01%) | 2643464059<br>(99.44%) | 2631343815<br>(98.99%) | 2280780358<br>(99.48%) | 2272198827<br>(99.11%) |
| No. of Mapped reads & % | 8245608<br>(40.6%) | 8133838<br>(40.0%) | 5829590<br>(26.6%) | 5737474<br>(26.1%) | 5121210<br>(19.5%) | 5046695<br>(19.2%) | 5422150<br>(23.9%) | 5353755<br>(23.6%) |
| No. of mapped reads have multiple alignment & % | 2896197<br>(35.1%) | 2848700<br>(35.0%) | 2122632<br>(36.4%) | 2084538<br>(36.3%) | 1890493<br>(36.9%) | 1858306<br>(36.8%) | 5422150<br>(23.9%) | 5353755<br>(23.6%) |
| No. Aligned pairs | 6510967 |  | 336496 |  | 3674315 |  | 3877244 |  |
| Total No. of expressed Genes | 33767 |  | 29510 |  | 32059 |  | 20152 |  |
|  | Low Zinc & Iron Wheat Genotypes |  |  |  |  |  |  |  |
|  | HUW-234 |  | AM-175 |  | AM-177 |  |  |  |
|  | HUW234_F | HUW234_R | AM175_F | AM175_R | AM177_F | AM177_R |  |  |
| No. of Raw reads | 23341420 | 23341420 | 22556866 | 22556866 | 19844961 | 19844961 |  |  |
| No. of HQ reads & | 23108745 | 23108745 | 22329403 | 22329403 | 19647224 | 19647224 |  |  |

| Percentage | (99%) | (99%) | (98.99%) | (98.99%) | (99.00%) | (99.00%) |
| --- | --- | --- | --- | --- | --- | --- |
| **No. of bases in HQ Reads** | 2333983245 | 2333983245 | 2255269703 | 2255269703 | 1984369624 | 1984369624 |
| **No. of HQ bases in HQ Reads** | 2321679049<br>(99.47%) | 2321679049<br>(99.02%) | 2243319127<br>(99.47%) | 2232331814<br>(98.98%) | 1973444626<br>(99.45%) | 1964578030<br>(99.00%) |
| **No. of Mapped reads & %** | 7213035<br>(31.2%) | 7108637<br>(30.8%) | 11289154<br>(50.6%) | 11133258<br>(49.9%) | 6687470<br>(33.7%) | 6549840<br>(33.0%) |
| **No. of mapped reads have multiple alignment & %** | 2446937<br>(33.9%) | 2404863<br>(33.8%) | 3875966<br>(34.3%) | 3809226<br>(34.2%) | 2225108<br>(33.3%) | 2167284<br>(33.1%) |
| **No. Aligned pairs** | 5489553 |  | 9209340 |  | 5065109 |  |
| **Total No. of expressed Genes** | 35785 |  | 36721 |  | 29369 |  |

### Functional annotation of DEGs, HSGs, and LSGs

Annotation of functions and description of DEGs, HSGs, and LSGs were identified by performing alignment with NR and UniProt database. The up-regulated DGEs **(Figure 3 & 4)** classified into heat shock proteins (HSPs), photosystem proteins, ABC transporter, ATP synthase family proteins, chloroplast protein and regulator of reverse DNA transcription proteins while the major classes of down-regulated genes are 30S/50S ribosomal chloroplast proteins, ATP-dependent Clp protease proteolytic subunit chloroplastic proteins, chlorophyll a-b binding of LHCII type protein, heavy metal transport proteins **(Supplementary Table S13 & S15)**. Similarly, annotations of HSGs (**Supplementary Table S17**) are classified into various genes and proteins families. HSGs families are 1-aminocyclopropane-1-carboxylate oxidase homolog 1-like family, ABC transporter family, acyltransferase family, aspartic family, ATP-dependent Clp protease family, accumulators, ATP-dependent zinc metalloprotease FTSH chloroplastic family, B-box zinc finger family protein, chloroplastic protein family, chaperone family, Choline ethanolamine kinase, calcium-binding family, cytochrome family, E3 ubiquitin family, ethylene-responsive transcription factor family, GPI ethanolamine phosphate transferase family. GTP-binding family, heat shock protein family, hypothetical proteins, phosphatidylinositol transferase, glutathione S-transferase family, a regulator of rDNA transcription, serine-threonine kinase family, transcription factors, metal binding, numerous transporters, and regulators. Likewise, LSGs are categorized by different genes and proteins families (**Supplementary Table S19)** [Gupta et al., 2018].

**Figure 3:**
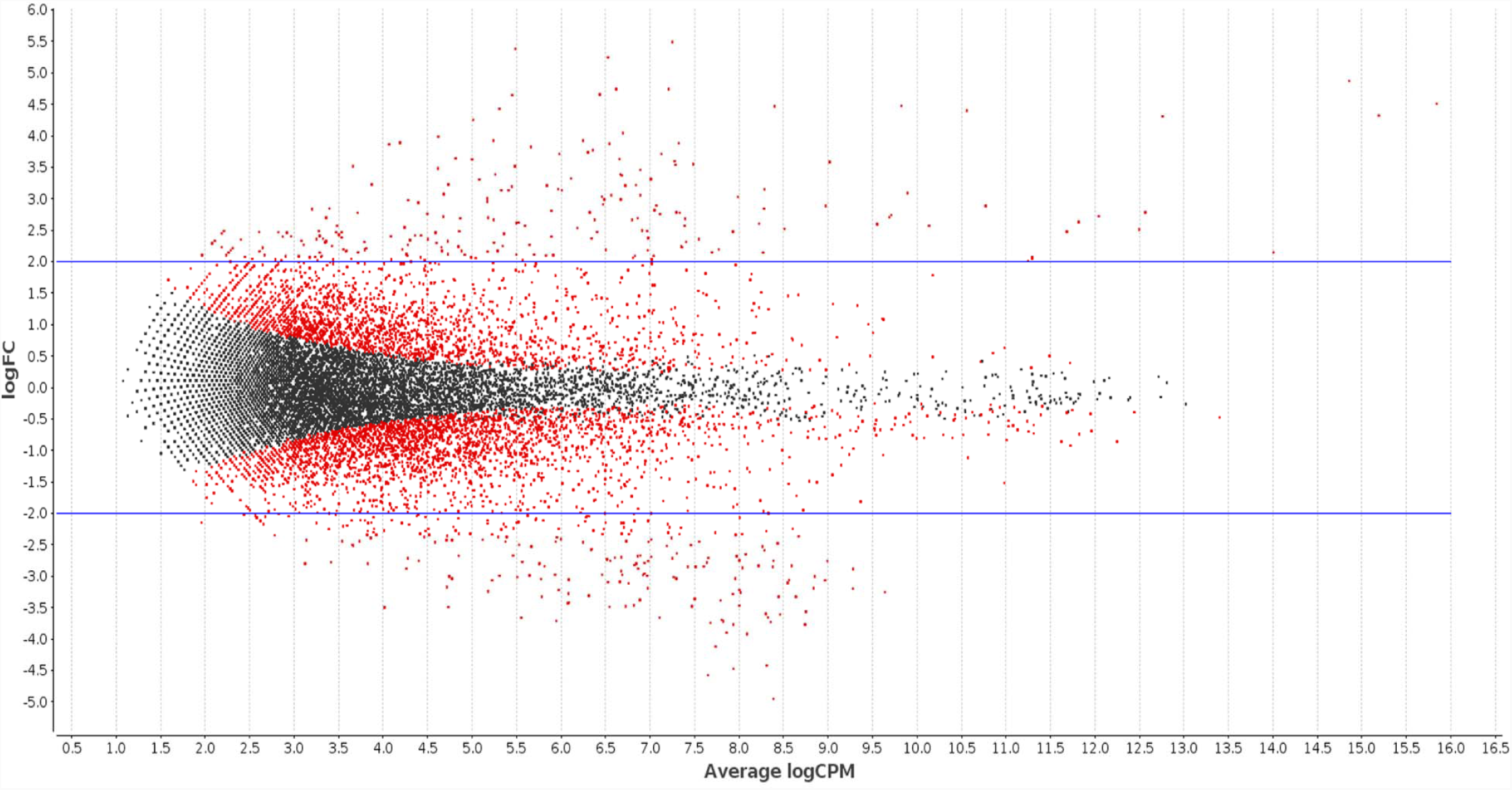
MA plot of differentially expressed genes identified in High and Low Zn & Fe accumulating Wheat Genotypes. Data represent individual gene responses plotted as logFC (Fold-Change) versus Average logCPM, with a negative change representing the down-regulated genes and a positive change representing the up-regulated genes. Black and Red points represent the genes having False Discovery Rate (FDR) >0.05 and < 0.05 respectively.

**Figure 4:**
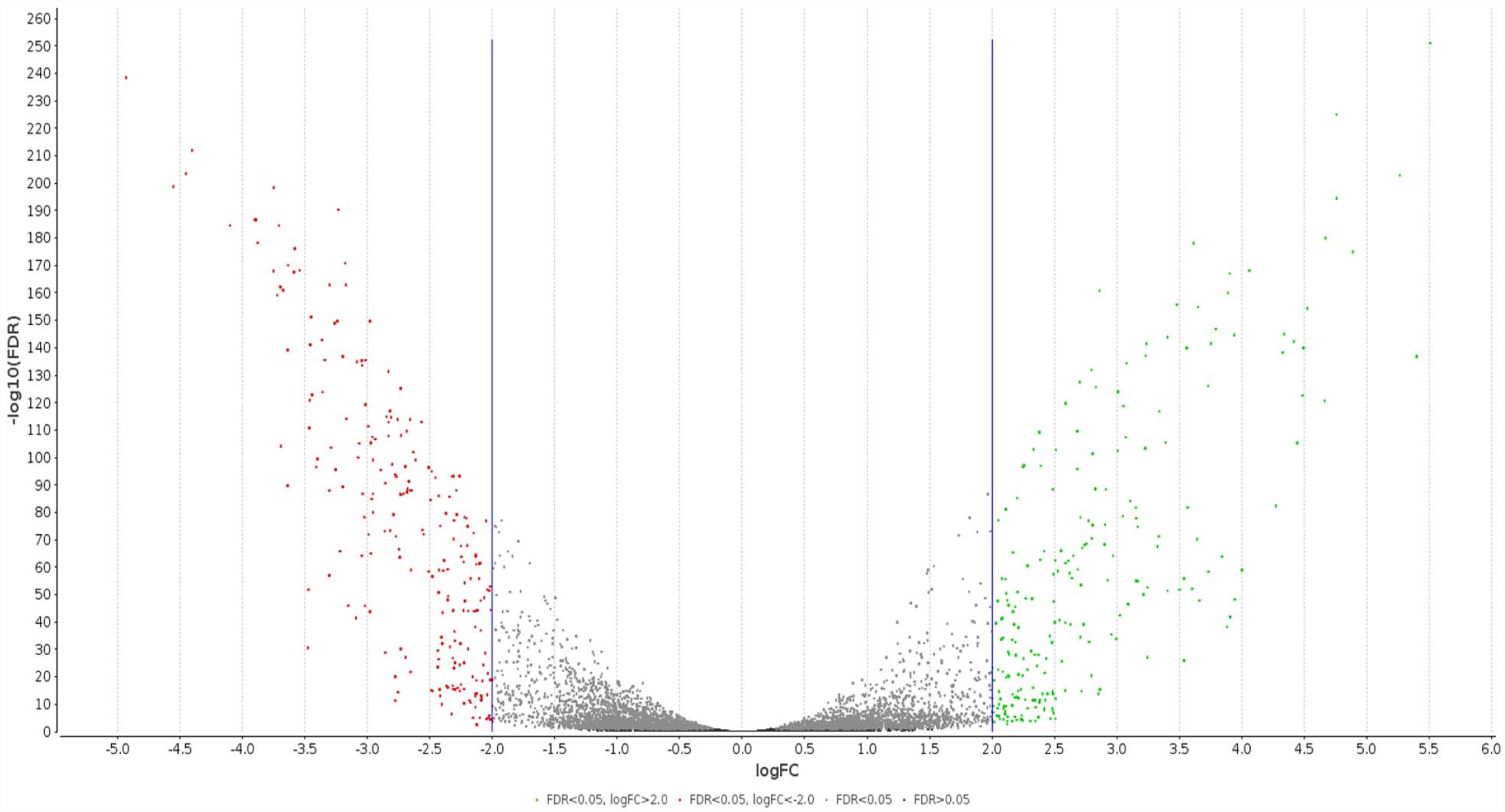
Volcano plot showing differentially expressed genes between high and low Zn & Fe accumulating wheat genotypes. The negative log10 transformed FDR values test the null hypothesis of no difference in expression levels between high Zn and low Zn wheat genotype (y axis) and are plotted against the average log fold changes (FC) in expression (x axis). Genes that were not classified as differentially expressed are plotted in black. In Red and Green are classified as differentially expressed and represents as down and up-regulated gens respectively.

The expression profiles of genes have a considerable difference between the two genotypes. To determine the functional significance of the transcriptional changes in each genotype, GO-based classifications were implemented for the up- and down-regulating genes as well as HSGs & LSGs. Only 212 up-regulated and 237 down-regulated genes were significantly enriched and assigned 877 and 1217 GO term respectively [Gupta et al., 2017a; Gupta et al. 2017b]. GO terms of biological process(BP) infers that oxidation-reduction process, protein-chromophore linkage, photosynthetic electron transport in photosystem, regulation of transcription, response to stress and stimulus resistance, ATP-Synthase coupled transport, protein and carbohydrate metabolism were found up-regulated **(Figure 5a)**, whereas GO terms translation, plastid translation, response to light stimulus, photosynthesis, proteolysis, chaperone-mediated protein folding mechanism are associated with down-regulated genes **(Figure 5d)**. In contrast, highly enriched BP indicates that most of the up-regulated enzymes (Photosynthetic enzymes, adenyl-pyrophosphatase, and nucleoside-triphosphate phosphatase) are involved in different metal ions accumulating metabolic processes [Varanavasiappan and Yeh, 2013]. The GO terms associated with various molecular functions (MFs), *viz.* ATP, chlorophyll Zn & Fe ion and other metal ions binding activity, electron transport, oxidoreductase activity etc. are highly enriched in up-regulated genes and support the aforesaid finding **(Figure 5b)**. However, MFs i.e. ribosomal formation and binding activities are highly enriched in the case of the down-regulated genes **(Figure 5e)**. Cellular components (CCs) such as an integral component of membranes, mitochondrion, nucleus, cytoplasm, chloroplast, plasma membrane, photosystem-II, plastids, ribosome and chloroplast **(Figure 5c & 5f)** found enriched, indicates their participation in the uptake of micronutrients. Zn & Fe are involved in chlorophyll synthesis, and also contributing to the maintenance of chloroplast structure and its function [Rout and Sahoo, 2015]. Similarly, GO analysis of top 450 LSGs and HSGs reveals that 375, 417 respectively, genes only enriched into distinguishing BP, MF, and CC GO terms. BP GO terms regulation of oxidation-reduction process, regulation of transcription & translation, synthesis of DNA template, protein phosphorylation, proteolysis and response to stress process are highly active in HSGs as compared to LSGs. The metal binding, Zn & Fe binding, and different cellular activities are highly active in HZFWGs as compared to LZFWGs corroborate the Zn & Fe accumulations facts of the wheat genotypes. Overall, this analysis provides differentiation in gene expression patterns between the HZFWGs and LZFWGs was corresponding to their distinct responses of Zn & Fe accumulation.

**Figure 5:**
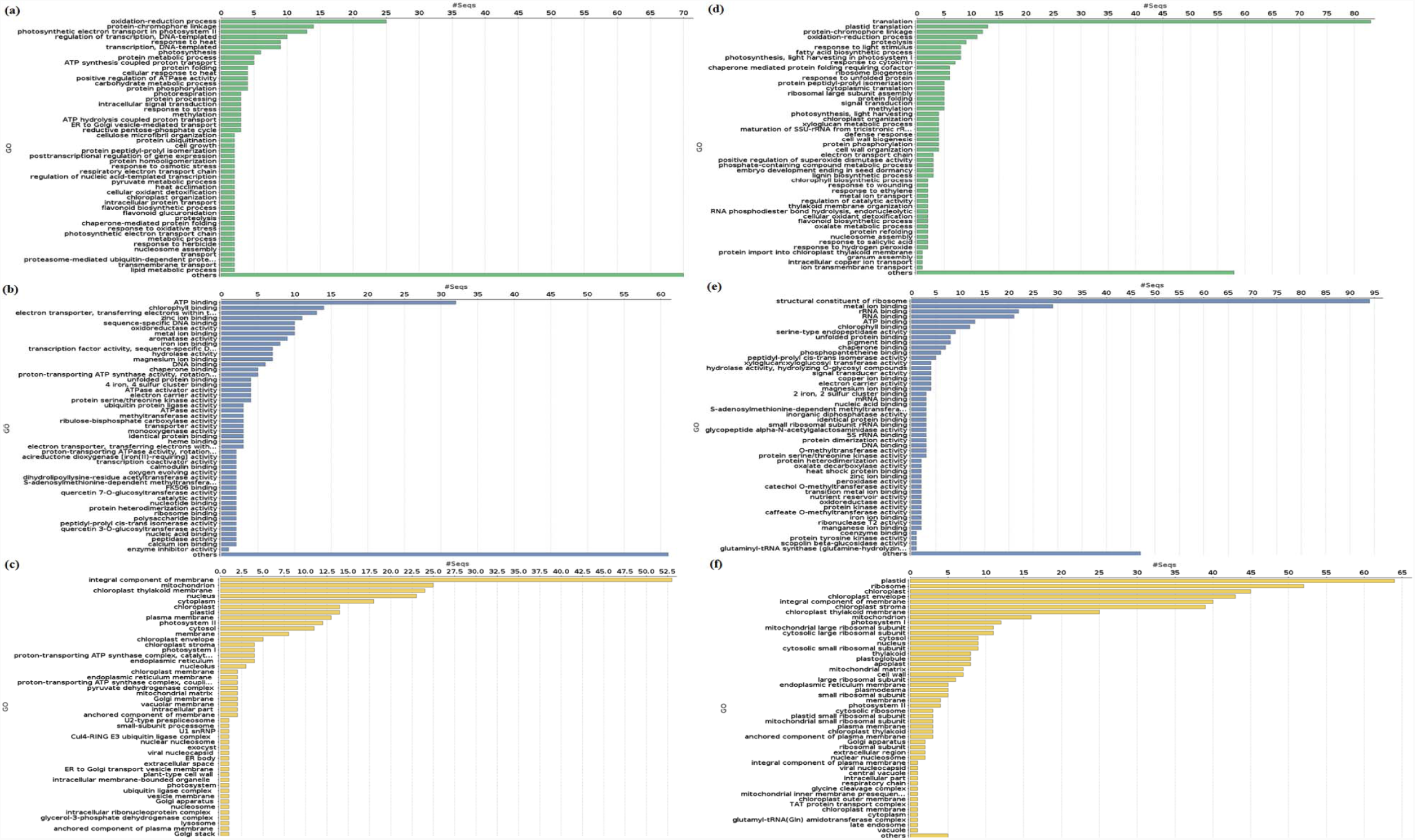
GO enrichment analysis of differentially expressed genes includes up- and down-regulated gene dataset. The total 449 DEGs were classified into (a) up-regulated- (d) down-regulated-biological process; (b) up-regulated- (e) down-regulated-molecular function; (c) up-regulated- (f) down-regulated-cellular component subcategories.

### HZFWGs and LZFWGs genes involvement in response to various stress

The transcriptomic analysis between HZFWGs and LZFWGs deduce that a higher number of transcripts was expressed in LZFWGs. However, considering the fact of genotype-specific gene expression also infers that LZFWGs expressed the higher number of specific genes. The HSGs and LSGs GO enrichment analysis revealed a significant difference between the two genotypes were observed in terms of enriched 1549 and 1306 GO terms respectively. The accumulation of minerals (Zn, Fe, Cu, Ca, Cu, Cd etc.) from soil to plant, is a complex regulatory process and directed by different metal ion binding activity and its transportation into different plant tissue. In contrast, the associated transcripts in Fe & Zn binding and transportation GO term found more in HZFWGs in comparison to LZFWGs **(Supplementary Table S17 & S19)**. GO terms were overrepresented in HSGs and categorized into BP, MF, and CC, especially those are related to stress tolerance *viz*. response to stress, response to biotic/abiotic stress, response to oxidative stress/osmotic stress, response to salt stress and response to heat. In HSGs 11 genes were involved in activating the response to stresses and encoding HSPs, dehydrin DHN3-like protein, LEA etc. [Gupta et al. 2016a]. Similarly, considering LSGs 16 genes are involved in response to stress, of these 5 genes associated with oxidative stress and 5 genes of salt stress indicating their high effects on LZFWGs. The major genes response to salt stress are S-adenosylmethionine synthase, BAG family molecular chaperone regulator 4 and Nitrogen regulatory P-II homolog and to oxidative stress named as tocopherol chloroplastic, Peroxidase A2, CBL-interacting protein kinase and chloroplastic lipocalin [Goldgur et al. 2007; Jin et al. 2016]. Likewise, 10 up-regulated genes are mainly encoding HSPs and associated proteins to respond to heat stress [Gupta et al., 2015]. These genes were playing an important role in protecting plant cells from environmental stress damage, by maintaining protein folding and regulation of other associated signaling proteins and metabolic pathways. It is noteworthy that without exposing any biotic/abiotic stress in the growth condition in these plants responding to stresses indicating their natural response phenomenon.

### Enhanced phenylpropanoid biosynthesis involved in STR in LZFWGs

To find out the differences in these metabolic processes took place in HZFWGs and LZFWGs the KEGG pathway enrichment analysis was performed in DEGs, HSGs, and LSGs. 33, 29, 58 and 63 KEGG pathways are enriched for up-regulated, down-regulated, HSGs and LSGs respectively shown in **Figure 6** and listed in **Supplementary Table S14, S16, S18 & S20**. Among all these pathways phenylpropanoid biosynthesis is highly enriched along with many other essential pathways such as glycolysis/gluconeogenesis, steroid hormone biosynthesis, pyruvate metabolisms are found common. In phenylpropanoid biosynthesis, 6 participatory enzymes are encoded by the genes of both genotypes **(Figure 7)**. It takes the input of phenylalanine/tryptophan biosynthesis pathway product and used intermediates enzymes to generate the secondary metabolites i.e. flavonoids, hydroxycinnamic acids, esters, guaiacyl, syringyl, and lignin. With the combination of reductases, oxygenases, and transferases, they are also amplified in numerous cascades to developed specific organ/tissues plant. Involved enzymes i.e. gentiobiase (EC:3.2.1.21), lactoperoxidase (EC:1.11. 1.7), dehydrogenase (EC:1.1.1.195), O-methyltransferase (EE: 2.1.1.68), ammonia-lyase (EC:4.3.1.24 and EC: 4.3.1.25) are encoded by down-regulated genes, however dehydrogenase, gentiobiase, and lactoperoxidase are commonly encoded by up-regulated HSGs and LSGs. This indicates LZFWGs possibly producing higher secondary metabolites such as flavonoid biosynthesis, common lignin, coumarins, phenolic volatiles, or hydrolyzable tannins etc. combinedly accountable to respond STR [Vogt 2010; Weisshaar and Jenkins 1998]. These results imply the phenylpropanoid biosynthesis pathway involvement defense response to STR in LZFGWs (Dixon et al., 1996), also affirmed from the GO enrichment analysis of LZFGWs associated expressed genes.

**Figure 6:**
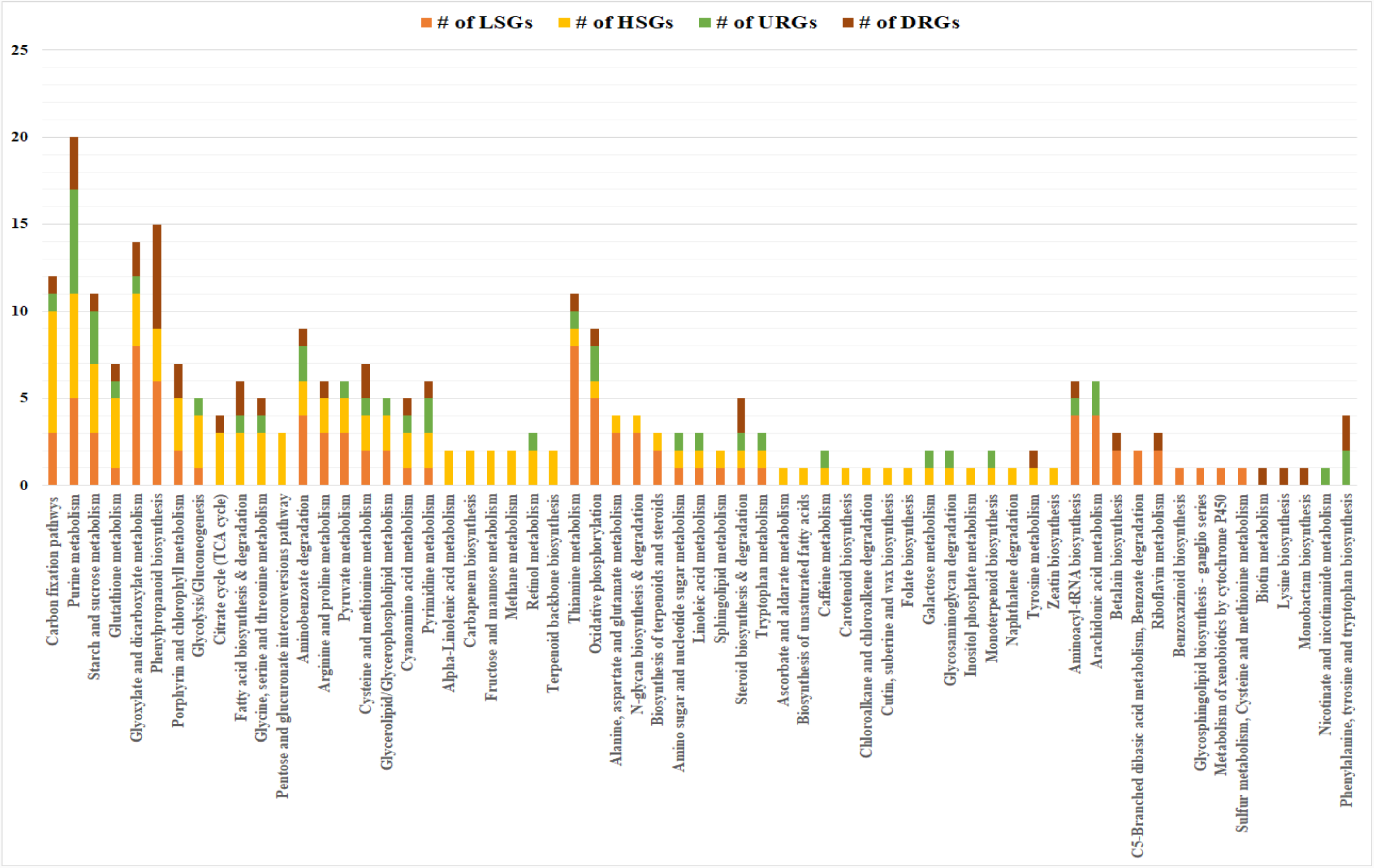
Bar graph indicates the metabolic pathways and associated gene number (#) belongs to different categories. Maximum number of enzymes from high and low specific genotypes are participated in different paathways.

**Figure 7:**
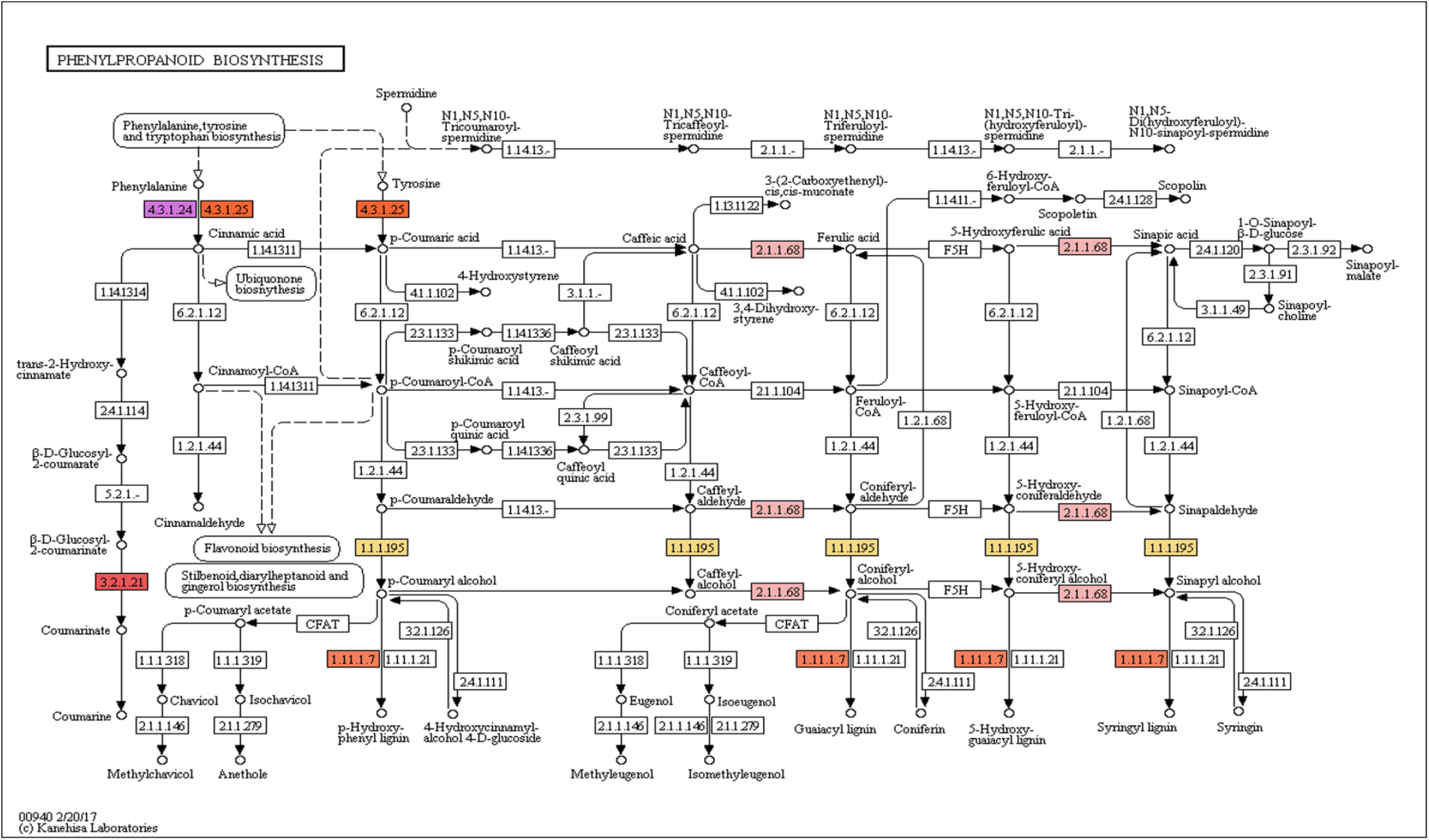
Pathway of the of Phenylpropaniod biosynthesis in evolved from KEGG. Several genes involved to encode the highlighted enzymes of this pathway, involved in plant stress response. The enzymes are Gentiobiase (EC: 3.2.1.21), Lactoperoxidase (EC: 1.11.1.7), Dehydrogenase (EC: 1.1.1.195), O-methyltransferase (EC: 2.1.1.68), Ammonia-lyase (EC: 4.3.1.24) and Ammonia-lyase (EC: 4.3.1.25) are induced inside the plant cell.

### Differentially expressed transcripts participating in uptakes of mineral transportations

The minerals absorbing processes in the plant from the soil is governed by different accumulators. Generally, they are translocated in the shoot and another part of the plant and transfer nutrients and minerals to their consumption’s location of the plant. Minerals uptake is affected by pH, water content, organic substances present in the soil. Moreover, minerals accumulation requires a suitable transporting system, regulated by different gene regulator and attributes to enhance the minerals uptake [Waters et al., 2009]. Various studies reported minerals accumulators also participates in response to stress tolerance or resistance (STR) [Kumar et al. 2018; Shabala et al., 2014; Moon et al., 2014]. We have identified such transcripts those are functioning as mineral accumulators as well as STR (Table 3). Number of proteins were expressed in our analysis, such as ABC transporter family proteins (4), calcium-dependent kinase family (1), chloroplastic family proteins (7), CBL-interacting kinase family (2), cytochrome family protein (2), glutathione peroxidase (4), helicase-like transcription factor family (1), mitogen-activated kinase (1), Na+ H+ antiporter(1), peroxidase family protein (4), serine-threonine kinase family (12), wall-associated receptor kinase family (7) and zinc finger transcription factor family (9) are participating in minerals accumulation as well as STR. Tow transcripts of heavy metal transport detoxification superfamily were expressed, that is involved in mineral accumulation and transportation only (Table 3). In addition, protein family such as abscisic stress ripening proteins (1), heat shock protein family (16), heat stress transcription factor (15), chloroplast enhancing stress protein (1) and WRKY transcription factors (1) are functionally involved in response to STR [Gupta et al 2018; Gupta et al 2019].

**Table 3:** List of transcripts having dynamic association with minerals uptake and stress tolerance and resistance.

| Transcript ID | Protein family | MAF | STR | Predicted Functions | Reference |
| --- | --- | --- | --- | --- | --- |
| TraesCS4D01G000500.1;<br>TraesCS4A01G000500.1<br>TraesCS2A01G110200.1;<br>TraesCS3A01G536300.1 | ABC transporter family proteins | Yes | Yes | ATP binding; ATPase activity, coupled to transmembrane movement of substances | Shabala et al., 2014;<br>Moon et al., 2014 |
| TraesCS4D01G109500.1 | Abscisic stress ripening | No | Yes | Response to stress | Goldgur et al., 2007 |
| TraesCS3A01G350700.1;<br>TraesCS3D01G344700.1 | BAG family molecular chaperone regulator | No | Yes | Response to stress; chaperone binding | Doukhanina et al., 2006 |
| TraesCS2D01G404200.1 | Calcium-dependent kinase family | Yes | Yes | Abscisic acid-activated signalling pathway; calcium ion binding; calmodulin-dependent protein kinase activity | Sharma et al., 2018 |
| TraesCS7A01G196500.1;<br>TraesCS2B01G496600.1<br>TraesCS2A01G541400.1;<br>TraesCS6B01G371500.1<br>TraesCS7D01G162300.2;<br>TraesCS6D01G320800.1<br>TraesCS5B01G321800.1; | Chloroplastic family proteins | Yes | Yes | Metal ion binding & transportation; oxidation-reduction process; response to stresses | Watson et al., 2018 |
| TraesCS2D01G097700.1;<br>TraesCS5D01G388000.1 | CBL-interacting kinase family | Yes | Yes | ATP binding; protein serine/threonine kinase activity; protein phosphorylation; intracellular signal transduction; response to stresses | Jin et al., 2016 |
| TraesCS1D01G183000.1;<br>TraesCS5D01G221600.1<br>TraesCSU01G269500.1; | Cytochrome family protein | Yes | Yes | Iron ion binding; oxidoreductase activity, acting on paired donors, with incorporation or reduction of molecular oxygen, NADPH as one donor; oxidation-reduction process; response to stress; | Castorina et al. 2018;<br>Hänsch et al.,2009 |
| TraesCS7B01G041000.1<br>TraesCS5A01G224100.1;<br>TraesCS3A01G458200.1<br>TraesCS3D01G084600.1;<br>TraesCS3A01G153000.1<br>TraesCS3B01G179500.1;<br>TraesCS1B01G347700.1<br>TraesCS5A01G318100.1;<br>TraesCS4A01G460100.1 |  |  |  |  |  |
| TraesCS1B01G128900.1;<br>TraesCS1D01G112400.1<br>TraesCS4A01G322600.1 | E3 ubiquitin- ligase family | No | Yes | Ubiquitin protein ligase activity; proteasome-mediated ubiquitin-dependent protein catabolic process; response to stress | Serrano et al.,2018 |
| TraesCS3B01G234100.1;<br>TraesCS4B01G331100.1<br>TraesCS3D01G130700.1;<br>TraesCS2B01G083800.1 | Glutathione peroxidase | Yes | Yes | Response to oxidative stress; regulation of root development; glutathione peroxidase activity; metal ions binding & transportation | Martinez et al.,2018 |
| TraesCS3B01G348200.1; | Helicase-like transcription factor family | Yes | Yes | ATP & Zinc ion binding and Transportation, negative regulation of mRNA splicing; response to stresses | Sarwat et al., 2018 |
| TraesCS4B01G243400.1;<br>TraesCS2B01G047400.1<br>TraesCS3D01G351900.1;<br>TraesCS4B01G205700.1<br>TraesCS7A01G270100.1;<br>TraesCS7D01G270600.1<br>TraesCS1D01G382900.1; | Heat shock protein family | No | Yes | Response to heat stress; response to osmotic stress | Perk et al., 2015;<br>Gupta et al., 2016 |
| TraesCS7B01G117600.1<br>TraesCS4B01G225400.1;<br>TraesCS4D01G226000.1<br>TraesCS5D01G492900.2;<br>TraesCS3D01G218800.1<br>TraesCS1D01G121000.1;<br>TraesCSU01G116400.1<br>TraesCS4D01G206600.1;<br>TraesCS4A01G068200.1 |  |  |  |  |  |
| TraesCS7B01G267300.1;<br>TraesCS7A01G360400.1<br>TraesCS5A01G237900.1;<br>TraesCS7D01G440200.1<br>TraesCS3D01G280600.1;<br>TraesCS4D01G276500.1<br>TraesCS4A01G027700.1;<br>TraesCS4B01G278100.1<br>TraesCS2B01G419600.1;<br>TraesCS2A01G401600.1<br>TraesCS2D01G399000.1;<br>TraesCS7A01G451100.1<br>TraesCS7B01G351000.1;<br>TraesCS7A01G451100.1<br>TraesCS7B01G351000.1 | Heat stress transcription factor | No | Yes | Transcription factor activity, sequence-specific DNA binding; DNA-templated; response to stresses | Nover et al., 1996 |
| TraesCS1D01G372600.1 | Mitogen-activated kinase | Yes | Yes | Metal ion transportation; Signal transducer, downstream of receptor, with serine/threonine kinase activity; response to stress | Sinha et al.,2011;<br>Kumar et al., 2018;<br>Wimalasekera et al., |
|  |  |  |  |  | 2018 |
| TraesCS2D01G123000.1 | Na<sup>+</sup> H<sup>+</sup> antiporter | Yes | Yes | Response to salt stress; sodium proton antiporter activity; sodium ion transmembrane transport; Vacuolar membrane; regulator of pH | Huang et al., 2018 |
| TraesCS1B01G270000.1;<br>TraesCS1D01G259500.1 | Heavy metal transport detoxification superfamily | Yes | No | Metal ion binding, metal ion transport, response to cytokinin | Dubey et al. 2018 |
| TraesCS2B01G346800.1;<br>TraesCS6A01G169100.1<br>TraesCS1B01G115900.1;<br>TraesCS1D01G096400.1 | Peroxidase family protein | Yes | Yes | Peroxidase activity; response to oxidative stress; metal ion binding; hydrogen peroxide catabolic process | Alzahrani et al. 2018 |
| TraesCS3A01G328300.1 | Chloroplast enhancing stress protein | No | Yes | Hyperosmotic salinity response; response to photooxidative stress; photosystem I assembly; response to water deprivation | Yokotani et al. 2010 |
| TraesCS6D01G223400.1;<br>TraesCS2A01G509600.1<br>TraesCS2D01G493700.1;<br>TraesCS2D01G577000.1<br>TraesCS3A01G230500.1;<br>TraesCS1B01G446500.1<br>TraesCS7A01G313600.1;<br>TraesCS5D01G450100.1<br>TraesCS5B01G163400.1;<br>TraesCS6B01G436400.1 | Serine threonine kinase family | Yes | Yes | Abscise acid-activated signalling pathway; protein serine/threonine kinase activity; protein phosphorylation; response to stresses | Mao et al.,2009<br>Shi et al., 2018 |
| TraesCS2B01G537500.1;<br>TraesCS6A01G411900.1 |  |  |  |  |  |
| TraesCS3B01G591700.1;<br>TraesCS3D01G527200.1<br>TraesCS5B01G495300.1;<br>TraesCS5B01G367300.1<br>TraesCS2B01G536200.1;<br>TraesCS2A01G582400.1<br>TraesCS5B01G035000.1 | Wall-associated<br>receptor kinase<br>family | Yes | Yes | Metal ions binding & transportation; cell surface<br>receptor signalling pathway; protein serine/threonine<br>kinase activity; protein phosphorylation; | Hou et al., 2005;<br>Kanneganti et al.,<br>2008 |
| TraesCS1A01G410500.1 | WRKY transcription<br>factors family | No | Yes | Transcription factor activity, sequence-specific DNA<br>binding; regulation of transcription; response to<br>stresses | Chen et al., 2014 |
| TraesCS3A01G093300.1;<br>TraesCS6B01G248000.1<br>TraesCS5D01G207400.1;<br>TraesCS5D01G490700.1<br>TraesCS6A01G099800.1;<br>TraesCS4D01G125700.1<br>TraesCS6D01G005100.1;<br>TraesCS5B01G491300.1<br>TraesCS5B01G490700.1 | Zinc finger<br>transcription factor<br>family | Yes | Yes | Transcription factor activity, sequence-specific DNA<br>binding; metal ion binding; transcription regulatory<br>region DNA binding; Stress response | Bouain et al., 2014;<br>Zang et al., 2016 |

**MAF:** Metal ions associated function **STR:** Stress tolerance and resistance

### Purine metabolism responds more in HZFWGs indicates the high minerals uptakes

Comprehensive information about the mechanics of minerals uptake/accumulation of from soil is available [Sperotto et al., 2018; Che et al. 2018]. In depth of metabolic process involved of Zn & Fe in wheat is limited, however from the GO analysis as well as literature it has been confirmed that they play significant role BPs such as photosynthesis, chlorophyll synthesis, respiration, nitrogen fixation, minerals uptake mechanisms, regulation of transporters in different environmental stress and DNA synthesis through the action of the ribonucleotide reductase [Kim and Rees, 1992; Reichard, 1993; Che et al. 2018]. They also work as an active co-factor of many enzymes which is essential for plant hormone synthesis *viz.* ethylene, lipoxygenase, abscisic acid etc. According to pathway enrichment analysis, the up-regulated and high specific genes of HZFWGs encoding enzymes are participating in the purine metabolism **(Figure 8)**. Pentose phosphate, thiamine, histidine and riboflavin metabolism are the major constituent of this metabolic pathway, responsible for biosynthesis of bio-molecules and minerals accumulation. 23, 14 overexpressed genes of up-regulated and HSGs respectively, encode the six key enzymes i.e. adenylpyrophosphatase (EC: 3.6.1.3), phosphatase (EC: 3.6.1.15), kinases (EC: 2.7.4.8 and EC: 2.7.4.3), RNA polymerase (EC: 2.7.7.6) and uridine 5’-nucleotidase (EC: 3.1.3.5). They are involved in numerous sub-pathways of pathways of purine metabolism i.e. cyclic-GMP/AMP, serine/threonine biosynthesis etc. **(Figure 8)**. Cyclic-GMP/AMP is essential for activation of intracellular protein kinases in response to the binding of a membrane-impermeable peptide, hormones, and minerals to the external cell surface. Serine/threonine is required for plant growth, developments and direct accumulators during minerals uptake [Ros et al., 2015; Gupta et al., 2015]. This implied that highly enriched purine metabolism and its sub-pathways, possibly increasing the Zn & Fe accumulations in HZFWGs, which was consistent with the observations obtained from the GO enrichment analysis of expression pattern.

**Figure 8:**
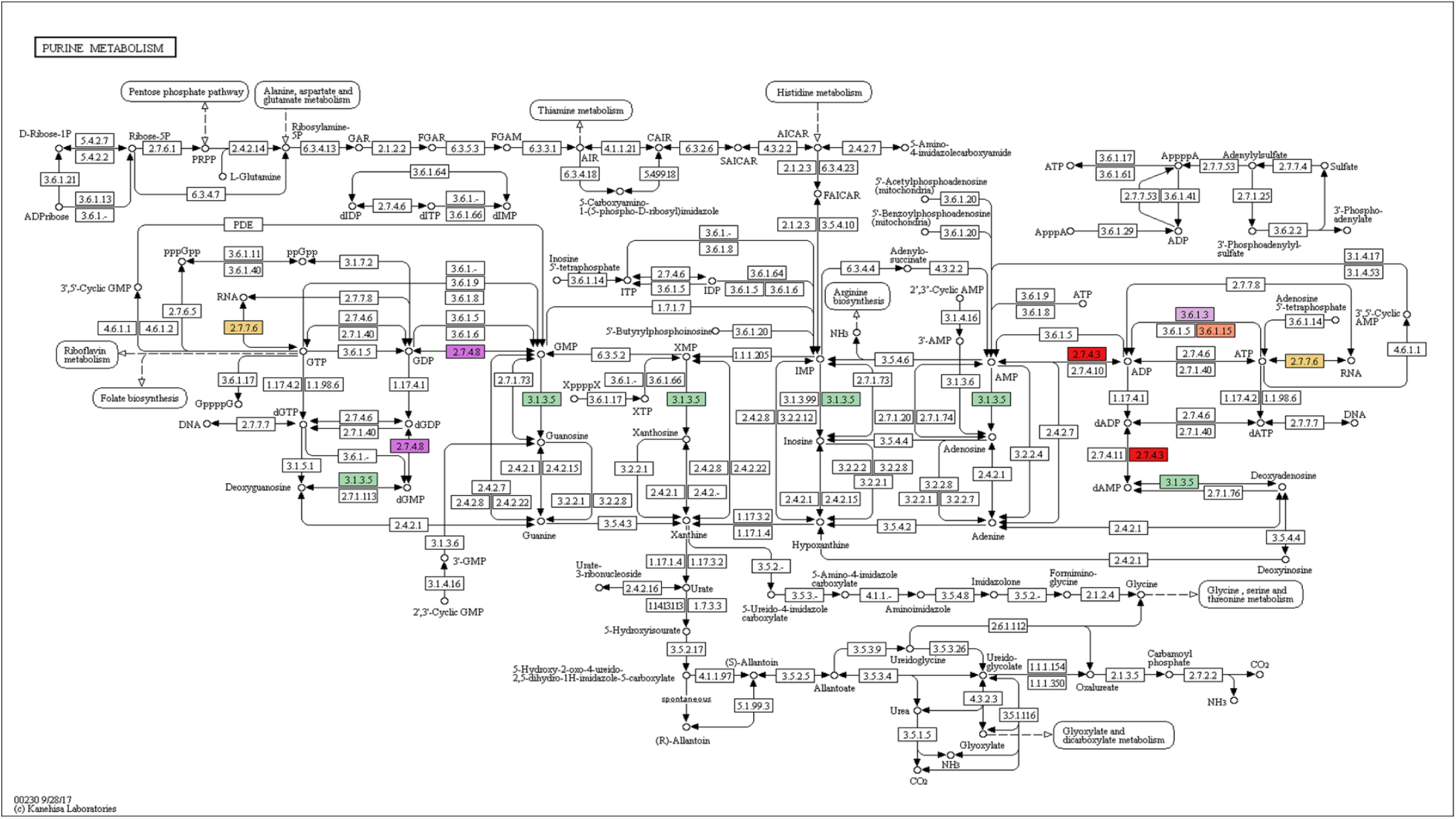
Pathway of the of Purine metabolism in evolved from KEGG. Several genes are involved to encode highlighted enzymes this pathway, to regulate plant growth and accumulation of minerals from soil. The enzymes Adenylpyrophosphatase (EC: 3.6.1.3), Phosphatase (EC: 3.6.1.15), Kinase (EC: 2.7.4.8), Kinase (EC:2.7.4.3), RNA polymerase (EC: 2.7.7.6), Uridine 5’-nucleotidase (EC:3.1.3.5) are participating to enhance the process in case of HZFWGs.

## Discussion

Globally spread malnutrition problem was mainly focused by various researchers across the world. The development the micronutrients rich cereal grains genotypes is one of the major constituents to overcome with this problem. Various efforts have been made to increase the concentration Zn & Fe in wheat through bioforctization [Cakmak, 2008; Velu et al. 2014; Velu et al. 2017; Khokhar et al., 2018; Cakmak, & Kutman, 2018] also studied the effects of drought & elevated temperature in grin Zn & Fe concentration [Velu et al. 2016]. Initially, QTLs map technique was widely used to obtained micronutrient rich genotypes [Velu et al. 2017]. Currently advance high through transcriptomic profiling analysis was preferred by researcher with respect OTL mapping to develop new plant genotypes [Panda et al. 2018; Zhou et al. 2016].

To determine the genes involved in minerals updates, numerous comparative transcriptomic studies have been performed in hyperaccumulating as well as in non-hyperaccumulating plant species [Gao et al. 2013; Di et al., 2011; Halimaa et al., 2014; Yang et al. 2017]. Current study also deals with comparative transcriptome analysis of 7 genotypes of low- and high- Zn & Fe accumulating wheat, contrasting the reason for grain mineral concentration by recognizing various DEGs associated in the uptakes of minerals (**Figure 1**). Results provide better understanding on metal homeostasis, metal tolerance, flavonoid biosynthesis, micronutrient, amino acid & protein transport and various stress responses in comparison earlier performed study by Singh et al. 2014 [Singh et al. 2014]. Prevalence of genes encoding proteins related Chlorophyll synthesis, Metal binding, Metal ion transport and ATP-Synthase coupled transport and stress responses, was noticed high in HZFWGs [Sperotto et al. 2014; Sperotto et al. 2018]. Furthermore, identified differentially expressed genes families (Table 3) found to be involved both Zn & Fe uptake and STR response in wheat. Identified ABC transporter and chloroplastic family proteins involved in ATP binding; ATPase activity metal ion binding and coupled to transmembrane movement of minerals [Shabala et al., 2014; Moon et al., 2014; Watson et al., 2018]. Expressed calcium-dependent kinase involved in abscisic acid-activated signalling pathway; calcium ion binding and its transportation; calmodulin-dependent protein kinase activity of wheat [Sharma et al., 2018]. Overexpression of cytochrome family proteins of wheat showing their accountability for metal ion binding & transportation; oxidation-reduction process and response to STR [Watson et al., 2018].

Number of proteins of CBL-interacting kinase, serine threonine kinase and wall-associated receptor kinase and Mitogen-activated kinase family were highly expressed wheat, involved in metal ion binding and its transportation serine/threonine kinase signalling, protein phosphorylation and response to stresses [Jin et al., 2016; Mao et al.,2009; Shi et al., 2018 Hou et al., 2005; Kanneganti et al., 2008; Kumar et al., 2018; Wimalasekera et al., 2018]. Glutathione peroxidase and peroxidase family proteins expression indicates higher response to oxidative stress, glutathione peroxidase activity, metal ions binding & transportation activity [Alzahrani et al. 2018; Martinez et al., 2018]. Interestingly, number of transcripts of zinc finger transcription factors and helicase-like transcription factor were expressed in HZFWGs showed their function in various transcription factor activity, sequence-specific DNA and metal ion binding and stress response [Bouain et al., 2014; Zang et al., 2016; Sarwat et al., 2018]. Accompanied number of transcripts of wheat are also expressed those are only involved in different STR. E3 ubiquitin-ligase family protein associated in proteasome-mediated ubiquitin-dependent protein catabolic process and response to stress [Serrano et al.,2018]. Heat shock protein family proteins and heat stress transcription factors overexpressed in both genotype indication their susceptible nature towards heat stress [Nover et al., 1996; Perk et al., 2015; Gupta et al., 2016].

Identified various metabolic process reveled their role in micronutrient uptake and stress (**Figure 6**). Exceedingly enriched phenylpropanoid biosynthesis metabolic pathways seemed to be participated in response to stress in LZFWGs were explored in the present study [Vogt, 2010]. Purine metabolism found to be contributing in the mineral uptakes in High- Zn & Fe accumulating wheat genotypes [Kim and Rees, 1992; Reichard, 1993]. These novel findings greatly enriched our knowledge on genetic basis of high- and low Zn & Fe accumulating wheat genotype. Further, this works illustrates the linkage between the different dimension of minerals accumulation and the various metal transition processes including Zn & Fe efflux, consumptions and translocation. These finding endow with some important facts can be used for developing High accumulating cultivars of wheat.

### Experimental Protocols

#### Plant genotypes and growth conditions

Total of 7 wheat genotypes were identified from Harvest Plus trials (4 HZFWGs and 3 LZFWGs) and validated in 20 villages for 2 years were selected for this experiment. Quantity of Zn and Fe concentration in seeds of each wheat genotypes was examined through a standard method (Ribeiro et al. 2012). High and low Zn & Fe concentration containing seed were sowed in twenty-one pots were filled with 30 kg homogenous soil (preparation given in next section) for growing 7 genotypes into 3 replications. This experiment was conducted during November to March 2015-16. Ten plants in each of pot were maintained at normal environmental conditions and grown up to ripening stage of plant. After 40 days of sowing at 12 leaf stage, 3 biological replicates of plant leaves of HZFWGs and LZFWGs were collated for total RNA extraction.

#### Preparation of homogeneous soil

To conduct the pot experiments, bulk surface soil sample (0-15cm) was collected from Agricultural Research Farm of Banaras Hindu University, Varanasi that was deficient in nutrients. The soil was air dried and ground to pass through 2 mm sieve. The soil samples were analyzed for pH, EC, organic carbon, available N, P, K, Zn, Fe, Cu, and Mn. Required quantities of fertilizers for 30 kg soil were calculated and applied in liquid form using Urea, DAP, Muraite of Potash and Gypsum as a source of N, P, K, and S respectively. The recommended N, P_2_O_5_ and K_2_O, S were 120, 60, 40 and 20 Kg ha^−1^ respectively. Half of N and full amount of P_2_O_5_ and K_2_O, S were applied at sowing. Remaining doses of nitrogen were added in 2 equal splits at 21 and 40 days after sowing. The experimental soil (0–15 cm) had pH 7.8 (1:2.5), EC 0.149 dS m^−1^, organic carbon 0.47% and available N, P and K of 130.2, 19.4 and 125.6 kg ha-1, respectively. The DTPA extractable Fe, Mn, Cu and Zn contents of soil were 42.5, 15.3, 2.4 and 0.51 mg kg^−1^, respectively. Samples of the soils were examined using standard procedures of by Sparks et.al (1996) [Sparks et al., 1996]. The DTPA extractable Fe, Cu, Mn, Zn, Cd, Cr, Ni and Pb were analyzed by atomic absorption spectrophotometer [Lindsay and Norwell, 1978].

#### RNA Extraction and High-Throughput Sequencing

Total RNA was extracted from 3 plant leaves as replicates of for each of LZFWGs (HUW-234, AM-175, & AM-177) and HZFWGs (BHU-3, BHU-35, BHU-22 & BHU-2) distinctly using standard Trizol RNA extraction protocol. The quality and quantity of RNA was confirmed by RNAse free agarose gel electrophoresis and concentration was measured using a 2100 Bioanalyzer [Agilent Technologies, Santa Clara, CA]. High-quality RNA was used for mRNA purification. Further, mRNA was purified using oligo-dT beads [TruSeq RNA Sample Preparation Kit, Illumina] using 1μg of intact total RNA. The purified mRNA was fragmented at 90°C in presence of divalent cations. The fragments were reverse transcribed using random hexamers and superscript II reverse transcriptase [Life Technologies]. Further, cDNA was synthesized on the first strand template using RNase and DNA polymerase-I and obtained cDNAs were cleaned using Beckman colter agencourt ampure XP SPRI beads. These cDNAs were ligated with Illumina adapters after end-repairing and the addition of an ‘A’ base followed by SPRI cleanup. The cDNA library was amplified using PCR for enrichment of the adapter-ligated fragments. The individual libraries were measured using a NanoDrop spectrophotometer and validated for quality with a Bioanalyzer. Subsequently, these libraries of each sample were used for RNA sequencing Illumina HiSeq 2500 platform [Illumina Inc., CA, USA]. The generated data files have been submitted to SRA database (Bio Project ID: PRJNA448363).

#### Preprocessing of RNA-Seq data and transcriptome profile analysis

Initially, the adapter sequences were removed from the raw reads. Then low-quality reads with having >50% bases with low-quality scores and/or >10% bases unknown (N bases) were removed from each dataset to gain more reliable results using NGSQC tool Kit [Patel et al., 2012]. The clean reads with the high-quality score of all samples were used in Top-Hat aligner [Kim et. al., 2013]. To perform the mapping with recently released high-quality reference transcriptome, assembled and released by International Wheat Genome Consortium in May 2017. The mapped reads of each sample were used for transcript assembly and quantification using StringTie assembler and Ballgown R package [Pertea et al., 2015; Frazee et al., 2015]. Assembler provides the annotated transcripts and their annotated length, Coverage (COV), Fragments per kilobase million (FPKM) and Transcripts per kilobase million (TPM) values for each sample. The genome-wide mapping of each expressed transcripts of all wheat genotypes sample were plotted in a circular plot using circos tool. This information of each sample was used to generate the 3 sets and 4 sets Venn diagrams for 3 LZFWGs and 4 HZFWGs respectively. This provides the commonly expressed transcripts among both types of genotypes. Moreover, common significant transcript among HZFWGs and LZFWGs were also identified. Further, differentially expressed genes (DEGs) between commonly expressed of HZFWGs and LZFWGs were calculated using Relative Log Expression (RLE) method of EdgeR Bioconductor package in R [Robinson et al., 2010].

#### Functional annotation GO and KEGG classification

Functional annotation of DEGs and top 450 of High Specific Genes (HSGs) and Low Specific Genes (LSGs) against databases Viz. NCBI non-redundant protein database (Nr), UniProt/Swiss-Prot database, InterPro Database, Kyoto Encyclopedia of Genes and Genomes (KEGG), by BLASTX searches with an e-value cutoff of 1e-5 in Blast2GO [Conesa et al.,2005]. To revalidate the annotation of the sequences BLASTX is used against specified Triticum aestivum databases of EnsemblePlants and Phytozome v12. Analysis provides multiple protein sequences for an input protein sequence, protein with the highest similarity score was considered as the optimal annotation and used for further study. GO classification and KEGG pathways analysis for HZFWGs as well as LZFWGs were performed for the up- & down-regulated genes and the chi-square test was employed to find out the GO terms of the significant difference in gene proportion between the two genotypes, which were proposed to play different roles in response Zn stress. For each KEGG pathway, the numbers of up- & down-regulated genes of each genotype were compared to the reference set by Fisher’s exact test to find out the pathways enriched with up and down-regulated genes. GO and KEGG enrichment analysis was also carried out for all the eight gene expression profiles.

#### SRA Database submission IDs

SRA database ID: SUB3858095 and Bio Project ID: PRJNA448363

## Acknowledgments

The authors would like to acknowledge Department of Applied Science of Indian Institute of Information Technology-Allahabad, India for providing the research infrastructure to carry out the whole transcriptomic data analysis.

## Contributions

SG, VKM, RC, AKJ, SKS, and RC conceived the work and designed the experiments. VKM, SG, and PKV performed all *In-silico* and *In-vitro* experiments. SG, VKM, RC, and PKV analyzed the results. SG, VKM, PKV, SKS, PSY, AKJ and RC contributed to writing the manuscript and discussed the results and commented on the manuscript.

## Compliance with Ethical Standards

This research does not perform any experiment on human and animals. Data used in this work were collected from open sources and all *In-vitro* data were generated at the Plant Genetic Lab, Banaras Hindu University-Varanasi India. Hence, the authors declare that there is no compliance with ethical standards.

## Legend for Supplementary Tables Information

Table S1: List of annotated transcripts of High Zn & Fe containing BHU-24 genotypes with their genomic location and different expression values.

Table S2: List of annotated transcripts of High Zn & Fe containing BHU-22 genotypes with their genomic location and different expression values.

Table S3: List of annotated transcripts of High Zn & Fe containing BHU-3 genotypes with their genomic location and different expression values.

Table S4: List of annotated transcripts of High Zn & Fe containing BHU-35 genotypes with their genomic location and different expression values.

Table S5: List of annotated transcripts of Low Zn & Fe containing AM-175 genotypes with their genomic location and different expression values.

Table S6: List of annotated transcripts of Low Zn & Fe containing AM-177 genotypes with their genomic location and different expression values.

Table S7: List of annotated transcripts of Low Zn & Fe containing HUW-234 genotypes with their genomic location and different expression values.

Table S8: List of commonly expressed transcripts in High Zn & Fe containing wheat genotypes with their genomic location and different expression values.

Table S9: List of commonly expressed transcripts of Low Zn & Fe containing wheat genotypes with their genomic location and different expression values.

Table S10: List of High Zn & Fe specific wheat genotypes expressed transcripts with their genomic location and different expression values.

Table S11: List of Low Zn & Fe specific wheat genotypes expressed transcripts with their genomic location and different expression values.

Table S12: List of up-, down-Normal-regulated transcripts of both types of genotypes with essential Fold change, FDR and P values.

Table S13: List of up-regulated genes significantly quantifying the GO terms with their gene description and other functional activities.

Table S14: KEGG enriched pathways list for up-regulated genes encoding different enzymes of each pathway.

Table S15: List of down-regulated genes significantly quantifying the GO terms with their gene description and other functional activities.

Table S16: KEGG enriched pathways list for down-regulated genes encoding different enzymes of each pathway.

Table S17: List of High Zn & Fe specific genes significantly quantifying the GO terms with their gene description and other functional activities.

Table S18: KEGG enriched pathways list for High Zn & Fe specific top 450 genes encoding different enzymes of each pathway.

Table S19: List of Low Zn & Fe specific genes significantly quantifying the GO terms with their gene description and other functional activities.

Table S20: KEGG enriched pathways list for Low Zn & Fe specific top 450 genes encoding different enzymes of each pathway.

## Supplementary Figure Information

**Figure S1:**
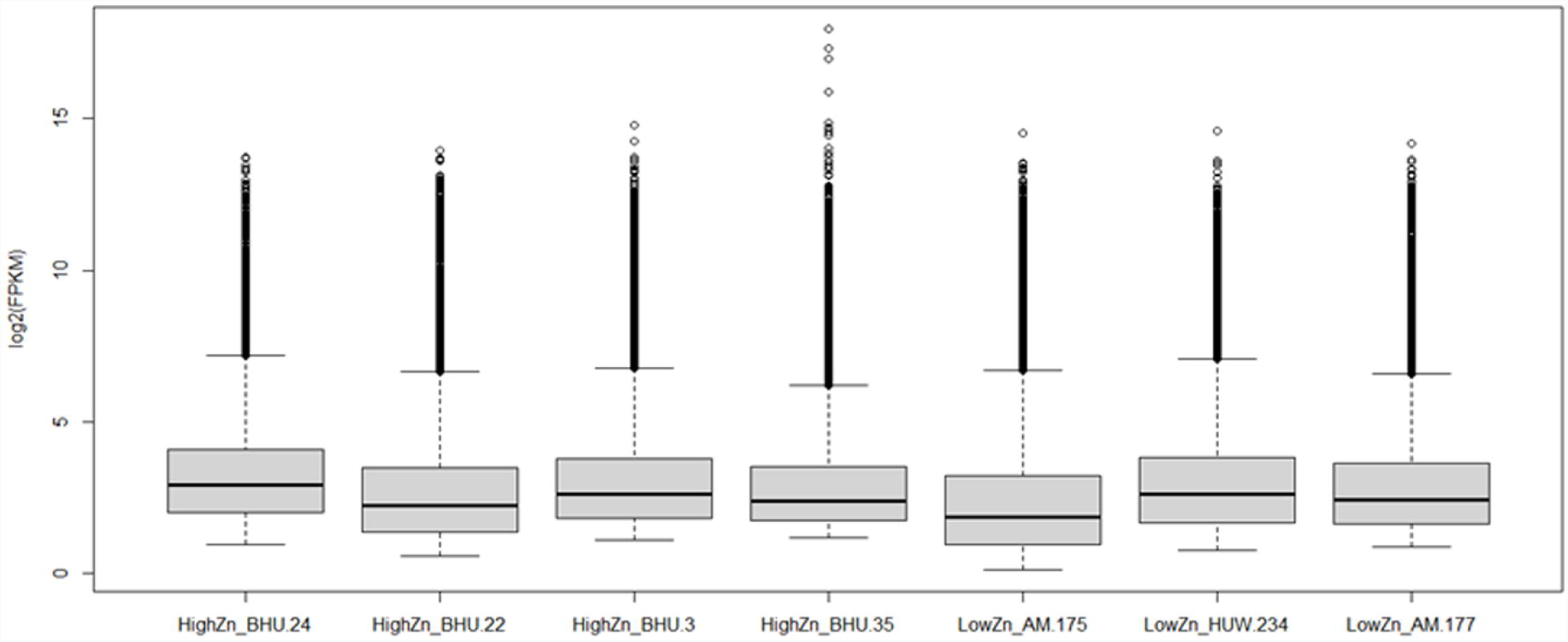
Box plot shows the level of gene expression [log2 (FPKM)] in High Zn & Fe and Low Zn & Fe accumulating wheat genotypes, denoted as HighZn_ & LowZn_ name of wheat genotype at the bottom of the plot.

**Figure S2:**
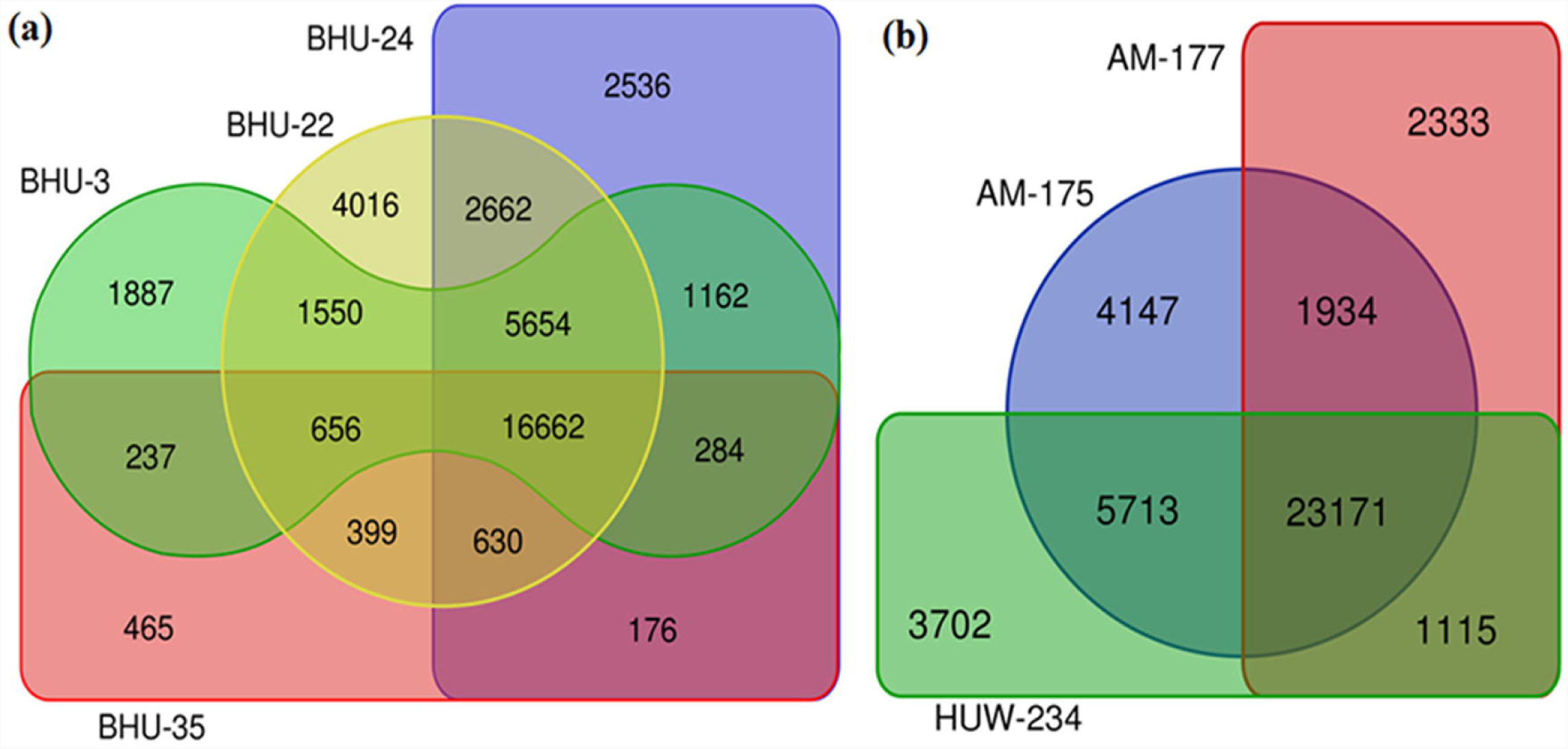
Venn diagrams show the common and unique annotated transcript among (a) High Zn & Fe-containing wheat genotypes (HZFWGs) and (b) Low Zn & Fe containing wheat genotypes (LZFWGs).

